# Integrated single-cell and bulk transcriptomic analysis leverages liver metastasis-related genes to develop a prognostic model for colorectal cancer patients

**DOI:** 10.64898/2026.03.24.714024

**Authors:** Xing Zhang, Wenxin Chen, Yalong Li, Lu Lu, Ruoming Huang, Jinglan Liao, Hong Li, Wanhui Zheng, Youqiong Xu

## Abstract

**Purpose:** Differentially expressed genes (DEGs) between colorectal cancer liver metastasis (CRLM) epithelium and primary colorectal cancer (CRC) epithelium (LMR DEGs) identified based on single-cell RNA sequencing (scRNA-seq) data may become new biomarkers for CRC prognosis.

**Methods:** An scRNA-seq dataset was used to describe the cellular landscape of primary CRC and CRLM and identify LMR DEGs. Prognostic LMR DEGs were identified in the bulk RNA-seq dataset. Based on the prognostic LMR DEGs, multiple machine learning algorithm combinations were compared in terms of their C-index, and the best model was selected for the construction of the LMR score.

**Results:** Among the 2070 LMR DEGs, 426 prognostic LMR DEGs were ultimately obtained. The combination of the randomized survival forest (RSF) model and ridge regression had the highest C-index and was therefore used to construct a 15-gene scoring system (LMR score). In the external validation set, the 1- and 5-year AUCs of the LMR score were greater than those of the AJCC stage and other scoring systems constructed with a similar dataset. In addition, the LMR score was closely associated with factors that influence CRC outcomes, such as immune infiltration.

**Conclusion:** The LMR score may be a reliable new biomarker for predicting the prognosis of patients with CRC.

## Introduction

Colorectal cancer (CRC) has one of the highest incidence and mortality rates of all malignant tumor types(1). The liver is the most common site of metastasis for CRC and colorectal cancer liver metastasis (CRLM) is the leading cause of death in patients with CRC(2, 3). However, the reasons for the tendency of CRC to metastasize to the liver are unclear, especially at the molecular level. The study of the mechanism of CRLM may contribute to clinical management and reduce mortality in patients with CRC. Recently, a study has noted the absence of colon gene programs in CRLM, which contrasts with the general view that metastatic tissue retains genetic features from the original tissue(4). This finding implies that in addition to anatomical proximity, genetic alterations may also contribute to the occurrence of CRLM. Therefore, an in-depth study of the differences in gene expression between CRLM and primary site CRC (primary CRC) may help explain the occurrence of CRLM and provide prognostic biomarkers for CRC patients.

Several studies based on bulk RNA sequencing (bulk RNA-seq) have focused on differentially expressed genes (DEGs) between CRLM and primary CRC (5–7). For example, Wang et al reported that IMPA2 could regulate CRLM by modulating lipid metabolism, epithelial mesenchymal transition (EMT) and DNA methylation; the expression of *IMPA2* has the potential to serve as a prognostic biomarker for CRC patients (7). These studies indicate the potential clinical value of aberrant gene expression in CRLM. However, the gene expression values obtained by bulk RNA-seq are affected by cells other than the epithelium, which may lead to inaccurate results. In recent years, single-cell RNA sequencing (scRNA-seq) technology has been rapidly developed and is capable of detecting gene expression at single-cell resolution(8, 9). The scRNA-seq technique allows for the targeted identification of DEGs between CRLM epithelial cells and primary CRC epithelial cells, and analysis of these DEGs may help us obtain more reliable biomarkers. Regrettably, no similar studies have been reported.

In our study, we first explored the cellular landscape of CRLM using a scRNA-seq dataset and CRLM-related DEGs (LMR DEGs) were identified by comparing the gene expression values between the CRLM epithelium and primary CRC epithelium. After that, we explored the prognostic predictive power of LMR DEGs and constructed a risk score system (the LMR score) based on prognostic LMR DEGs via an analysis of bulk RNA-seq datasets. Finally, we evaluated the associations between the LMR score and prognostic factors such as immune infiltration. With this study, we hope to improve our understanding of the development of CRLM and provide reliable biomarkers for the prognosis of CRC patients.

## Methods

### Acquisition of data

Supplementary Figure 1 shows the analysis flow of our study. The scRNA-seq data for 12 samples including 6 primary CRC samples and 6 CRLM samples, were downloaded from the GSE178318 dataset. Seven bulk RNA-seq datasets from the Cancer Genome Atlas (TCGA) database (TCGA-COAD and TCGA-READ) and Gene Expression Omnibus (GEO) database (GSE17536, GSE17537, GSE38832, GSE39582 and GSE161158) were downloaded for subsequent analysis. Survival information and clinical characteristics (age, sex, American Joint Committee on Cancer (AJCC) stage) of CRC patients were also collected. All the bulk RNA-seq datasets included disease-free survival (DFS) as the outcome variable, and patients with less than 1 month of follow-up were excluded. GSE161158 was used as an external validation set, and the remaining datasets were merged into the CRC meta-dataset. The “sva” R package is used to minimize batch effects between different datasets. In addition, methylation data of CRC patients in the TCGA database were also downloaded to explore the association between promoter region methylation levels and gene expression. The basic information of all the datasets included in this study is shown in Supplement Table 1.

### Quality control and analysis of the scRNA-seq data

Quality control and analysis of the scRNA-seq data were performed using the “seurat” R package. Cells with fewer than 500 genes detected (nFeature_RNA<500) and greater than 15% mitochondrial gene content (percent.mt>15%) were excluded. In addition, every gene must be detected in at least 3 cells. T-distributed stochastic neighbor embedding (t-SNE) was used for dimensionality reduction and visualization of the scRNA-seq data. Annotation information for cell clusters was derived from the “singleR” R package. The copy number variation (CNV) score was calculated for all epithelial cells using the inferCNV algorithm (based on the “inferCNV” R package) to identify malignant epithelial cells. In accordance with Zheng et al. (10), we defined malignant epithelial cells identified in primary CRC tissue as primary CRC epithelium and those identified in CRLM tissue as CRLM epithelium. Cell communication analysis for all cell types was performed via the “CellChat” R package. The FindAllMarkers() function was used to find the highly expressed marker genes for each cell type.

### Identification and enrichment analysis of DEGs

The FindAllMarkers() function was used to identify DEGs between primary CRC epithelium and CRLM epithelium, i.e., LMR DEGs (log fold change (logFC) >1, Bonferroni adjusted *P* <0.05). Gene set variation analysis (GSVA) analysis based on the “GSVA” R package was performed on the LMR DEGs. Prognostic LMR DEGs were identified using a multivariate Cox model (adjusted for age, sex and AJCC stage). Kyoto Encyclopedia of Genes and Genomes (KEGG) and Gene Ontology (GO) analyses were performed on the prognostic LMR DEGs using the “clusterProfiler” R package.

### Construction and validation of the LMR score and an online nomogram

The patients in the CRC meta-dataset were randomly divided into a training set and a test set at a 1:1 ratio. Referring to Liu et al. (11), 51 machine learning algorithm combinations were performed on the prognostic LMR DEGs to fit prediction models based on the basis of the leave-one-out cross-validation (LOOCV) framework in the training set. Machine learning algorithms include ridge, random survival forest (RSF), elastic network (Enet), least absolute shrinkage and selection operator (Lasso), CoxBoost, stepwise Cox, supervised principal component (SuperPC), partial least squares regression for Cox (plsRcox), survival support vector machine (survival-SVM) and generalized boosted regression model (GBM). The machine learning algorithm combination with the highest average C-index in the testing set and the validation set was considered to be the best predictive model for the prognosis of CRC patients. Based on the best prediction model, the LMR score is constructed as follows:

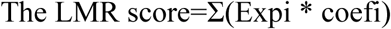

Coefi and Expi represent the coefficients and expression of the genes in the best prediction model, respectively. Patients were categorized into high- and low-risk groups according to the median LMR score. Gene set enrichment analysis (GSEA) was performed on DEGs (obtained via the “limma” R package) between the high- and low-risk groups using the “ReactomePA” R package. Kaplan‒Meier survival curves were used to compare the difference in survival between patients in the two groups. To evaluate the ability of the LMR score to predict DFS in CRC patients, we performed time-dependent receiver operating characteristic (t-ROC) curve analysis using the “survivalROC” R package to calculate the area under the curve (AUC). In the GSE161158 dataset, we compared the predictive efficacy of the LMR score with that of the pyroptosis-related gene (PRG) score (12), the cuproptosis-related gene (CRG) score (13) and the AJCC stage by calculating the 1-, 3- and 5-year AUCs. With the “rms” R package, a nomogram scoring system was constructed using the LMR score as well as clinical characteristics related to patient prognosis. Decision curve analysis (DCA) was conducted using the “dcurvesR” package to evaluate the clinical utility of the nomogram score. We used the “DynNom” R package to upload the nomogram scoring system to a web page for use by clinicians.

### Analysis of the key genes associated with the LMR score

The LMR DEG with the largest absolute value of the coefficient in the LMR score was evaluated for its ability to predict the prognosis of CRC patients using t-ROC curves. Mutations in this gene were searched for in the cBioPortal database and the association between its expression and promoter region methylation level was explored in the TCGA database.

### Immune infiltration analysis

The CIBERSORT algorithm was used to calculate the proportions of 22 immune-infiltrating cell types in the tissue. In the Tumor Immune Dysfunction and Exclusion (TIDE) database, we obtained TIDE scores, immune dysfunction scores, immune exclusion scores and response rates to immunotherapy for patients in the high- and low-risk groups. In addition, the expression of 33 common immune checkpoints was compared between the two groups.

### Assessment of chemotherapeutic drug sensitivity, tumor mutational burden and the cancer stem cell index

The “pRRophetic” R package was used to assess the sensitivity of CRC patients to commonly used chemotherapeutic agents. Tumor mutation burden (TMB) analysis and visualization were performed via the “mafTools” R package. The cancer stem cell (CSC) index was calculated for each patient in the TCGA dataset. The CpG island methylator phenotype (CIMP) index was derived for each patient in the TCGA dataset by calculating the percentage of CpG islands with an average methylation β-value ≥0.3; for each island, the methylation level was determined by averaging the β-values of all probes annotated to that CpG island(14, 15).

### Statistics

All the statistical analyses were conducted using the R software (version 4.1.1). Unless otherwise stated, all the statistical *P* values are two-sided and P<0.05 represents statistical significance.

## Results

### Cellular landscape of primary CRC and LM CRC

The t-SNE reduction divides 99,340 QC-eligible cells into 13 cell clusters, which further are annotated into 8 cell types (Supplementary Figure 2A-C, Figure 1A). InferCNV analysis was performed to identify malignant epithelial cells. Not all epithelial cells presented significant copy number variation (Figure 1B). The CNV scores for reference and epithelial cells were calculated and then clustering analysis was performed. We observed that cells in Class 1 had the lowest CNV score and consisted predominantly of reference cells (Figure 1C). Therefore, the epithelial cells in Class 1 were defined as normal epithelial cells (n= 1296) and the epithelial cells in other classes were defined as malignant epithelial cells. We defined malignant epithelial cells from primary CRC samples as primary CRC epithelium (n= 6879) and malignant epithelial cells from LM CRC samples as LM CRC epithelium (n=2247). All cell types and cell counts are shown in Supplementary Table 2. The distributions of the three epithelium types were close but clearly differentiated on the t-SNE plot, indicating similarity and heterogeneity in the transcriptional characteristics of the different epithelium types (Figure 1D). Cell communication analysis revealed that tissue stem cells closely interact with all three epithelium types, especially with the primary CRC epithelium and LM CRC epithelium (Figure 1E). The heatmap shows the top 5 marker genes for each cell type (Figure 1F).

**Figure 1.**
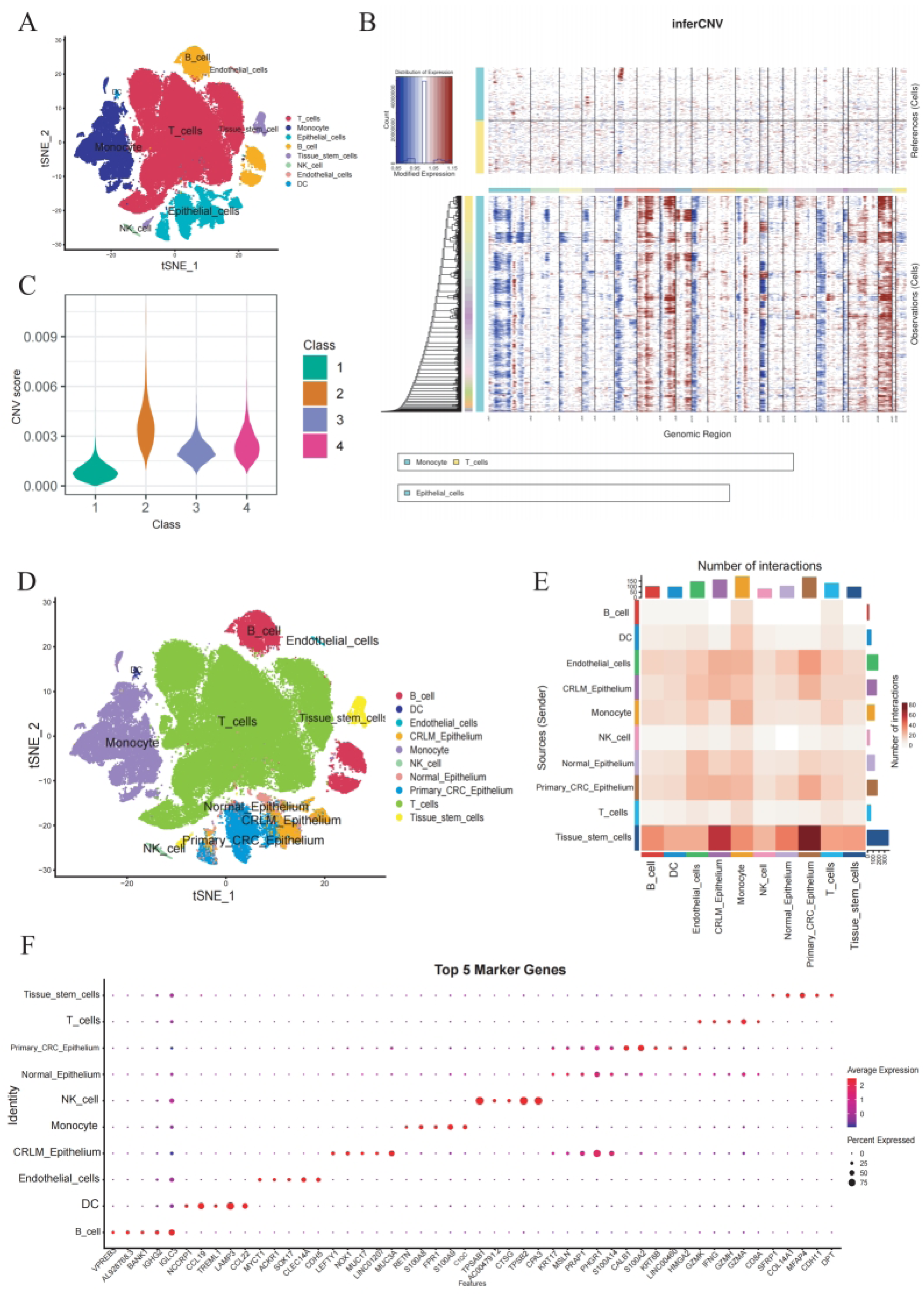
The t-SNE plots demonstrated cell types before (A) and after (D) epithelial cell subdivision. The heatmap (B) and violin (C) plot showed CNV status of epithelial cells. (E) Interactions between different cell types were explored by cell communication analysis. (F) The dotplot showed top 5 marker genes for all cell types.

### Identification of prognostic LMR DEGs

A total of 2070 LMR DEGs were identified in the CRLM epithelium compared with the primary CRC epithelium, including 1308 high-expressed LMR DEGs and 762 low-expressed LMR DEGs (Supplementary Table 3). GSVA of the LMR DEGs revealed that the PPAR signaling pathway, retinol metabolism, drug metabolism cytochrome P45, arachidonic acid metabolism, glycolysis gluconeogenesis, and lysosome pathways were significantly upregulated in the CRLM epithelium (Figure 2A). Next, we merged 6 bulk RNA-seq datasets (TCGA-COAD, TCGA-READ, GSE17536, GSE17537, GSE38832 and GSE39582) to form the CRC meta-dataset. In the CRC meta-dataset, we separately included each LMR DEG in a multivariate Cox regression model (adjusted for age, sex, and AJCC stage) to identify prognostic LMR DEGs. Finally, we obtained 426 prognostic LMR DEGs, including 283 risk prognostic LMR-DEGs and 143 protective prognostic LMR DEGs (Supplementary Table 4). KEGG analysis revealed that the prognostic LMR DEGs were mainly enriched mainly in focal adhesion, amoebiasis, cytokine−cytokine receptor interaction, human papillomavirus infection and ECM−receptor interaction (Figure 2B). GO analysis revealed that prognostic LMR DEGs were enriched in terms related to signaling (Figure 2C).

**Figure 2.**
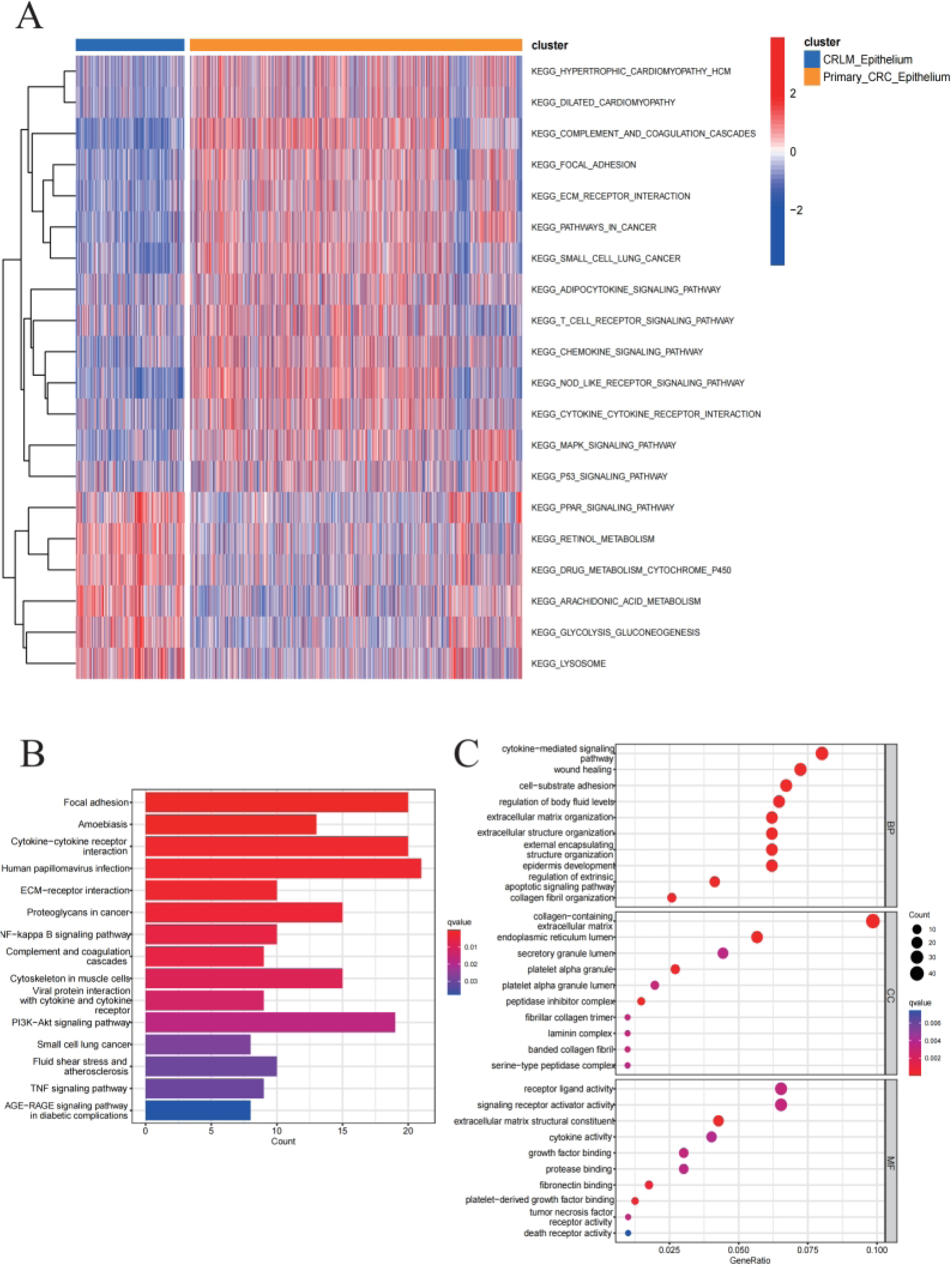
(A) GSVA analysis revealed abnormal up- and down-regulated pathways in CRLM epithelium. The KEGG (B) and GO (C) pathway enrichment analysis were performed on LMR DEGs.

### Construction and validation of the LMR score

The CRC meta-dataset was randomly divided into a training set and a testing set at a ratio of 1:1. The GSE161158 dataset was used as an external validation set. In the training set, 51 machine learning models based on prognostic LMR DEGs were produced to predict the DFS of CRC patients (Figure 3A). The combination of RSF and ridge regression had the highest average C-index and was therefore used to create the LMR score:

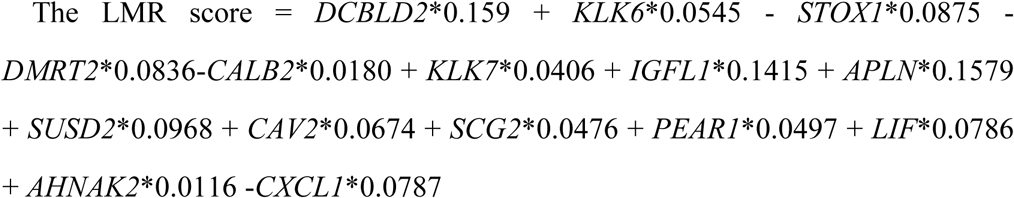

**Figure 3.**
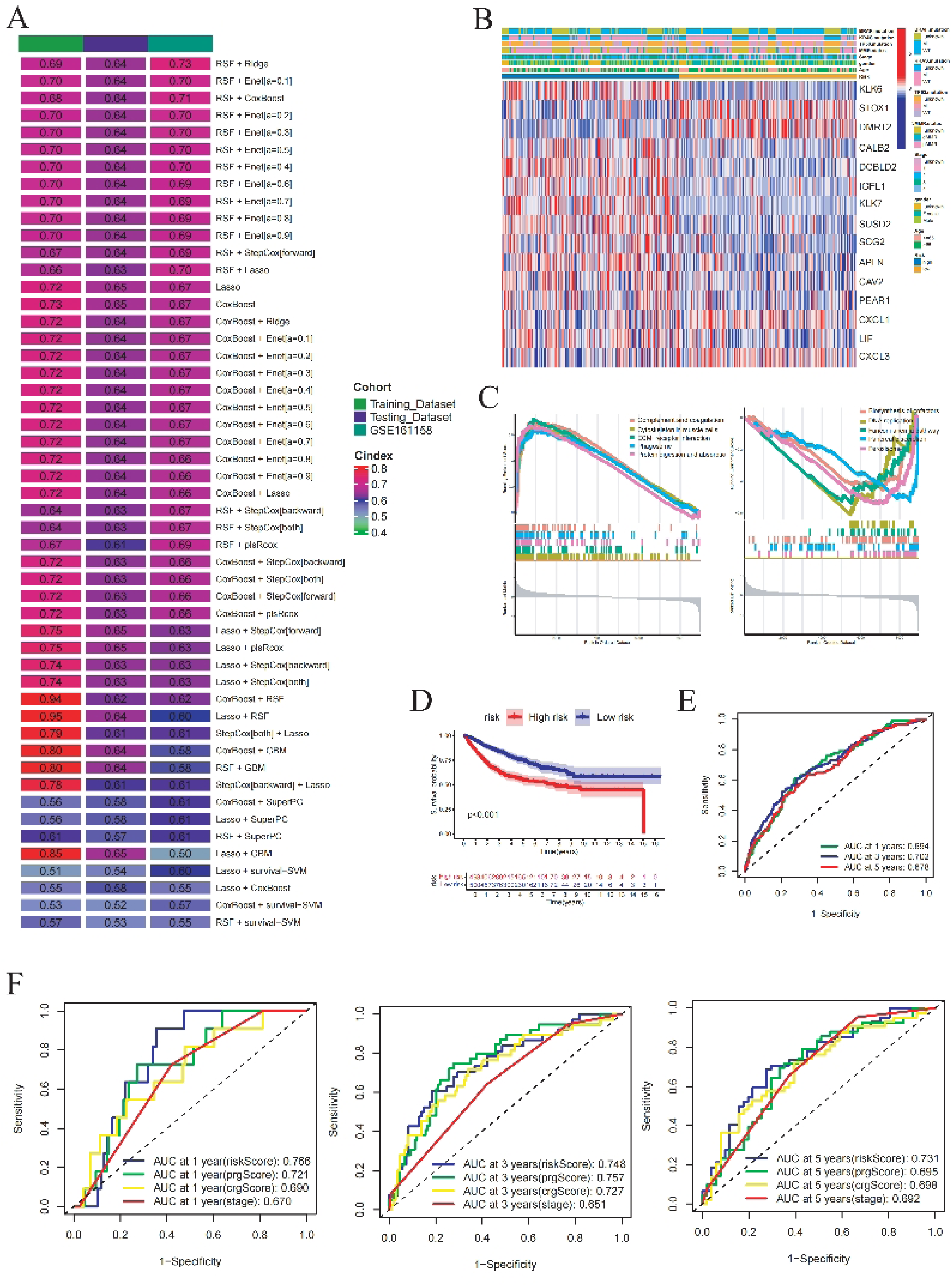
(A) C-index for a total of 51 machine learning algorithm combinations in the training set, testing set and validation set. (B) The heatmap showed the distribution of clinical features between high- and low- risk groups. (C) GSEA analysis revealed dysregulated pathways between high and low risk groups. The Kaplan-Meier survival curve (D) and the t-ROC curve (E) demonstrated the LMR score’s ability to differentiate the prognosis of CRC patients in the CRC meta-dataset. (F) The predictive effect of the LMR score, the PRG score, the CRG score and the AJCC stage on the prognosis of CRC patients was compared in the external validation set (GSE161158) using 1-, 3-and 5-year AUCs.

We calculated the LMR score for each patient. Patients were categorized into the high-risk group if their scores were above the median LMR score; otherwise, they were categorized into the low-risk group. The heatmap demonstrated the distribution of clinical features between the two risk groups (Figure 3B). GSEA of the DEGs between the two groups revealed that the five most significantly upregulated pathways in the high-risk group were complement and coagulation cascades, cytoskeleton in muscle cells, ECM-receptor interaction, phagosome and protein digestion and absorption, and the five most significantly downregulated pathways were biosynthesis of cofactors, DNA replication, the Fanconi anemia pathway, pancreatic secretion and peroxisomes (Figure 3C). Patients in the low-risk group had a significantly more favorable prognosis than those in the high-risk group did in all datasets (Figure 3D, Supplementary Figure 3A-C). Multivariate Cox regression revealed that the LMR score was an independent prognostic predictor for CRC patients (Table 1). The AUCs of the LMR score in the CRC meta-dataset were 0.694, 0.702, and 0.678, respectively (Figure 3E). The LMR score also had a similar predictive effect on both the training set and the testing set (Supplementary Figure 3D-E). We then compared the AUCs of the LMR score with thosse of the PRG score, the CRG score and the AJCC stage in the validation set. The 1-, 3-, and 5-year AUCs of the LMR score in the GSE161158 dataset were 0.766, 0.748, and 0.731, respectively. The LMR score had the highest AUC for 1- and 5-year DFS, and the AUC for 3-year DFS was slightly lower than that of the PRG score but higher than those of the AJCC stage and the CRG score (Figure 3F).

**Table 1:** Multivariate Cox regression analysis of CRC patients in the CRC meta-dataset and the validation set.

| Variables | CRC meta-dataset(N=998) |  |  |  | GSE161158 dataset (N=188) |  |  |  |
| --- | --- | --- | --- | --- | --- | --- | --- | --- |
|  | 95%CI |  |  |  | 95%CI |  |  |  |
|  | HR | Lower | Upper | P | HR | Lower | Upper | P |
| Age | 1.027 | 1.017 | 1.037 | <0.001 | 0.998 | 0.976 | 1.020 | 0.847 |
| Stage |  |  |  |  |  |  |  |  |
| II vs. I | 1.789 | 1.026 | 3.119 | 0.040 | 2.605 | 0.582 | 11.650 | 0.210 |
| III vs. I | 2.544 | 1.458 | 4.439 | 0.001 | 4.533 | 1.058 | 19.410 | 0.042 |
| IV vs. I | 7.524 | 3.947 | 14.342 | <0.001 | 5.307 | 0.786 | 35.840 | 0.087 |
| Gender |  |  |  |  |  |  |  |  |
| Male vs. Female | 1.592 | 1.260 | 2.010 | <0.001 | - | - | - | - |
| Risk score | 2.618 | 2.117 | 3.237 | <0.001 | 8.747 | 3.481 | 21.980 | <0.001 |

### Prognostic predictive effectiveness and expression regulators of *DCBLD2*

In the process of constructing the LMR score, the expression of *DCBLD2* had the largest coefficient, indicating that it plays an important role in predicting patient prognosis. The median *DCBLD2* expression value was used to categorize patients in the dataset into high- and low-expression groups. High *DCBLD2* expression was unfavorable for the prognosis of CRC patients (Figure 4A-C). The t-ROC curves revealed that the expression level of *DCBLD2* had a certain predictive ability for the DFS of CRC patients, regardless of the dataset (Figure 4D-F). We explored the association between the methylation and expression of *DCBLD2* in the TCGA database. The results revealed a significant negative correlation between the promoter region methylation level of *DCBLD2* and its expression level (Figure 4G). Analysis of the cBioPortal database revealed that the mutation rate of *DCBLD2* in CRC patients was 2.28% (Figure 4H). Mutations in *DCBLD2* were predominantly missense and involved three structural domains: CUB, LCCL and F5_F8_type_C (Figure 4I).

**Figure 4.**
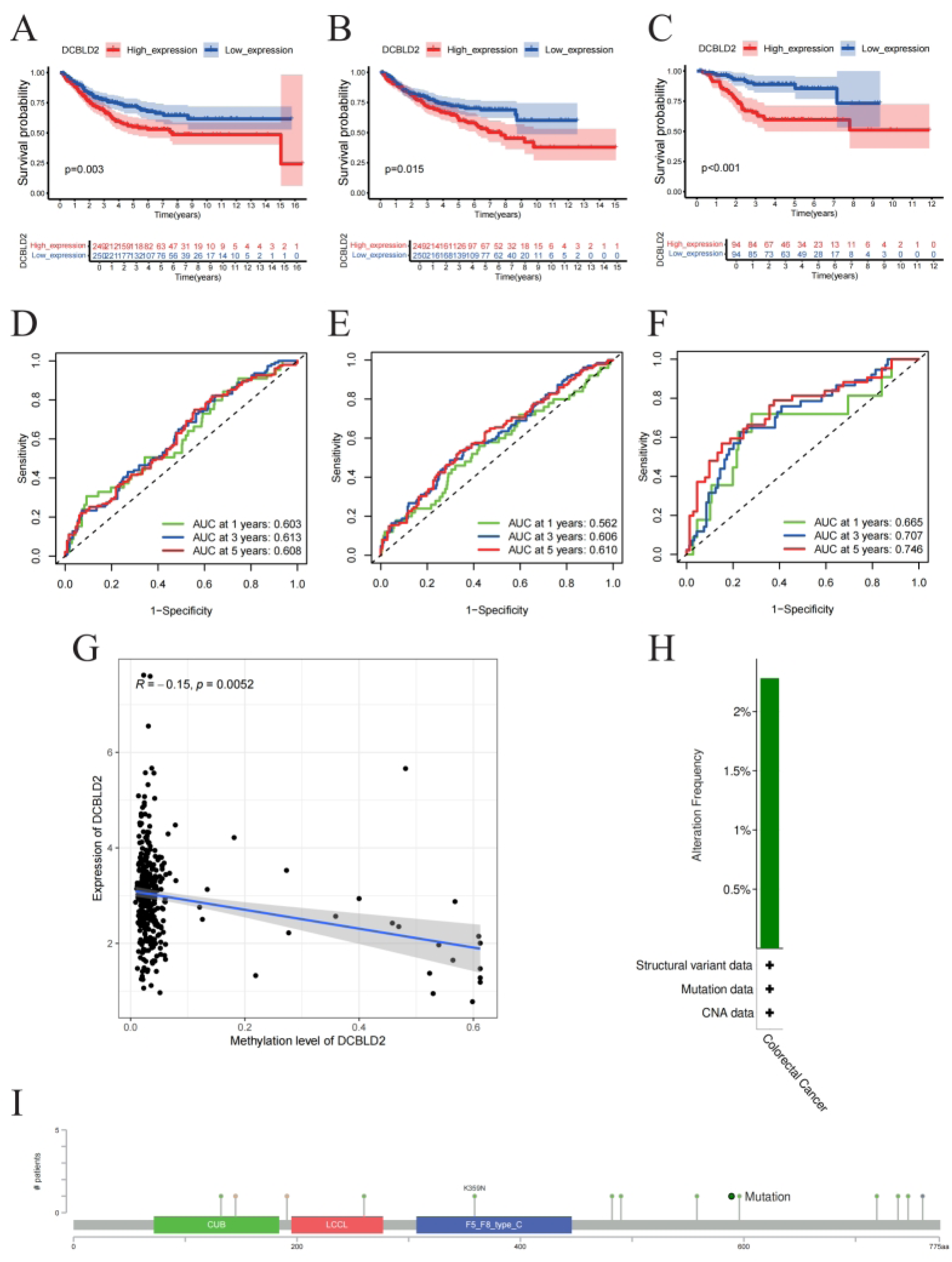
The Kaplan-Meier survival curve showed the difference in CRC patients’ survival between high and low *DCBLD2* expression groups in the training (A), testing (B) and validation sets (C). The t-ROC curves demonstrated the predictive ability of the expression of *DCBLD2* for the prognosis of CRC patients in the training (D), testing (E) and testing sets (F). (G)Association between promoter region methylation level and gene expression of *DCBLD2*. The bar plot (H) and the lollipop plot(I) showed mutation status of *DCBLD2* in the cBioPortal database.

### Construction of the nomogram scoring system

A nomogram scoring system was constructed on the basis of sex, age, AJCC stage, and the LMR score (Figure 5A). Patients whose nomogram score was higher than the median nomogram score were categorized into the high-risk group and the remaining patients were categorized into the low-risk group. Patients in the high-risk group had significantly worse prognoses than those in the low-risk group did (Figure 5B). The calibration curve revealed that the nomogram score had a high accuracy in predicting survival outcome (Figure 5C). The 1-, 3- and 5-year AUCs of the nomogram score were 0.716, 0.739 and 0.743, respectively, which were greater than those of the LMR score (Figure 5D). DCA demonstrated that the nomogram score provided a favorable net clinical benefit across a wide range of risk thresholds (Supplementary Figure 4A-C). A web-based version of the nomogram model was subsequently created for ease of clinical application (Figure 5E, https://zxwyy210822.shinyapps.io/Nomo/).

**Figure 5.**
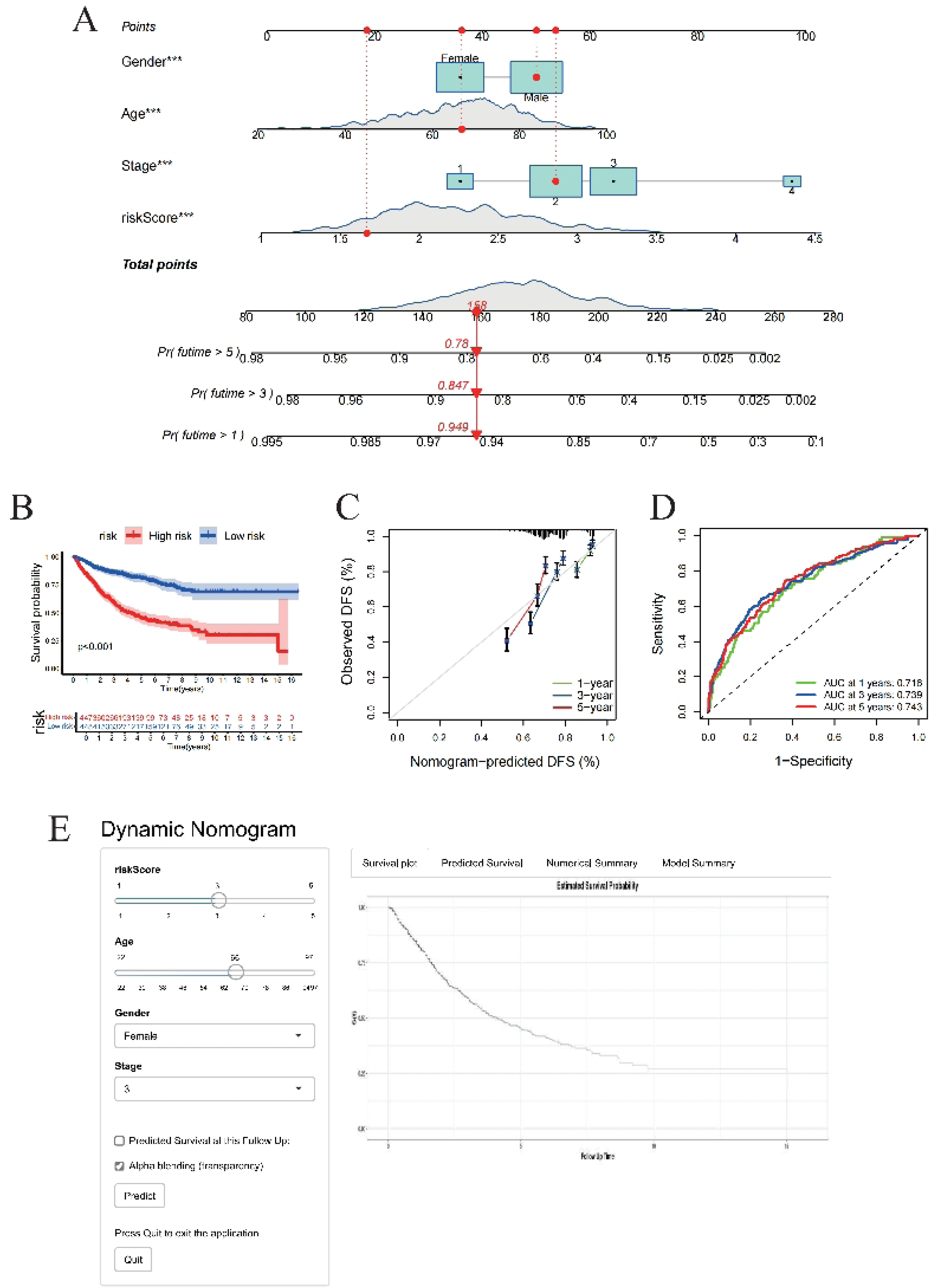
Construction of the nomogram scoring system. **(A)** The nomogram scoring system was constructed based on age, AJCC stage and the risk score. **(B)**The Kaplan-Meier survival curve showed the difference in CRC patients’ survival between high and low risk groups in the CRC meta-dataset. (C)The calibration curve used to evaluate the accuracy of the nomogram model’s predictions in the CRC meta-dataset. (D)The t-ROC curve showed 1-, 3- and 5-year AUCs for the nomogram model in the CRC meta-dataset. (E) An online prognostic prediction tool built based on our nomogram scoring system.

### Correlation of The LMR Score with Immune Infiltration

Next, we explored the correlation between the LMR score and tumor immune infiltration. All 15 genes used to construct the LMR score were associated with the abundance of immune cells, especially neutrophils (Figure 6A). The LMR score was negatively correlated with the numbers of resting NK cells, Tregs, plasma cells, activated memory CD4+ T cells, resting memory CD4+ T cells and naive B cells and positively correlated with the numbers of neutrophils, M0 macrophages and M2 macrophages (Figure 6B). The analysis of the TIDE database revealed that patients in the high-risk group had higher TIDE scores, immune dysfunction scores and immune exclusion scores than patients in the low-risk group did, whereas patients in the low-risk group had higher MSI scores (Figure 6C-F). Patients in the low-risk group had a greater response rate to immunotherapy (56.8% vs. 33.5%, Figure 6G). In addition, 19 of the 33 common immune checkpoints were significantly differentially expressed between the high- and low-risk groups (Figure 6H). These results indicated that patients in the low-risk group had better outcomes with immunotherapy.

**Figure 6.**
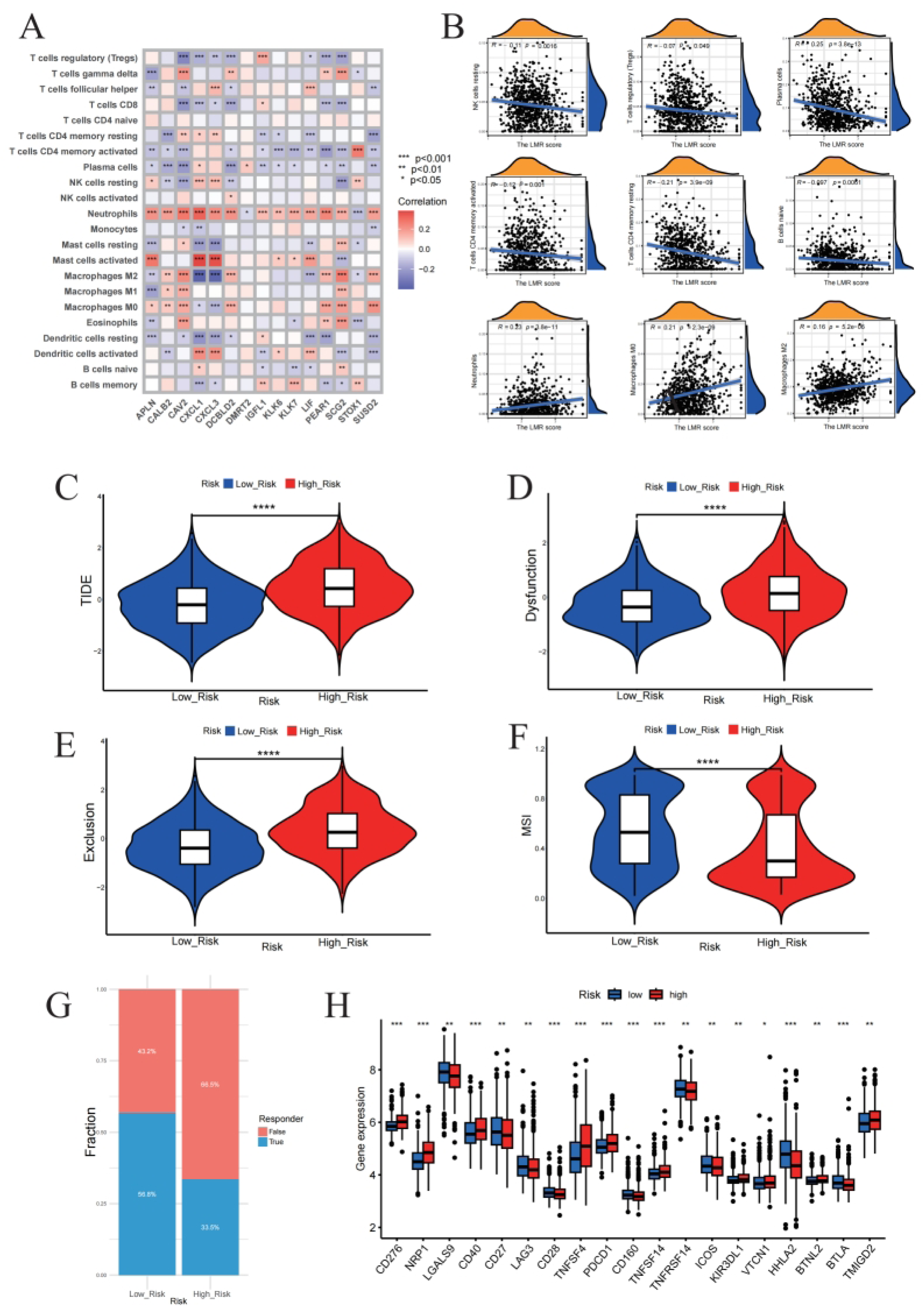
The correlation between the risk score and tumor immune infiltration. (A) The heatmap showed the correlation between the 15 LMR DEGs used to construct the LMR score and 22 immune cell types. (B)The plots demonstrated the immune cell types that were significantly associated with the LMR score. The Violin plots revealed the differences between the TIDE score (C), the dysfunction score (D), the exclusion score (E), and the MSI score (F) between high and low risk groups. (G) The bar plot showed the immunotherapy response rate in the high and low risk groups. (H) The box plot showed the difference in expression of 33 common immune checkpoints between the high and low risk groups.

### Correlation of the LMR Score with the TMB, CSC index, CIMP index and chemotherapy drug sensitivity

The waterfall plots revealed that the highly mutated genes were essentially the same in the high- and low-risk groups (Figure 7A). As RAS/BRAF mutation status is associated with adverse prognosis of CRC patients (16, 17), we specifically examined the mutation profiles of *KRAS*, *NRAS* and *BRAF* in the high- and low-risk groups. The results revealed a higher mutation rate of these three genes in the high-risk group (Supplementary Figure 5A-B). The LMR score showed no significant correlation with either TMB (R = 0.067, *P* > 0.05; Figure 7B) or the CIMP index (R = 0.061, *P* > 0.05; Supplementary Figure 5C). In addition, we found a significant negative correlation between the LMR score and the CSC index (R = − 0.35, *P* < 0.001; Figure 7C). Next, we explored the association between the LMR score and chemotherapeutic drug sensitivity in patients with CRC. We found that gefitinib, paclitaxel and shikonin had lower IC50 values in the high-risk group whereas gemcitabine had lower IC50 values in the low-risk group (Figure 7D).

**Figure 7.**
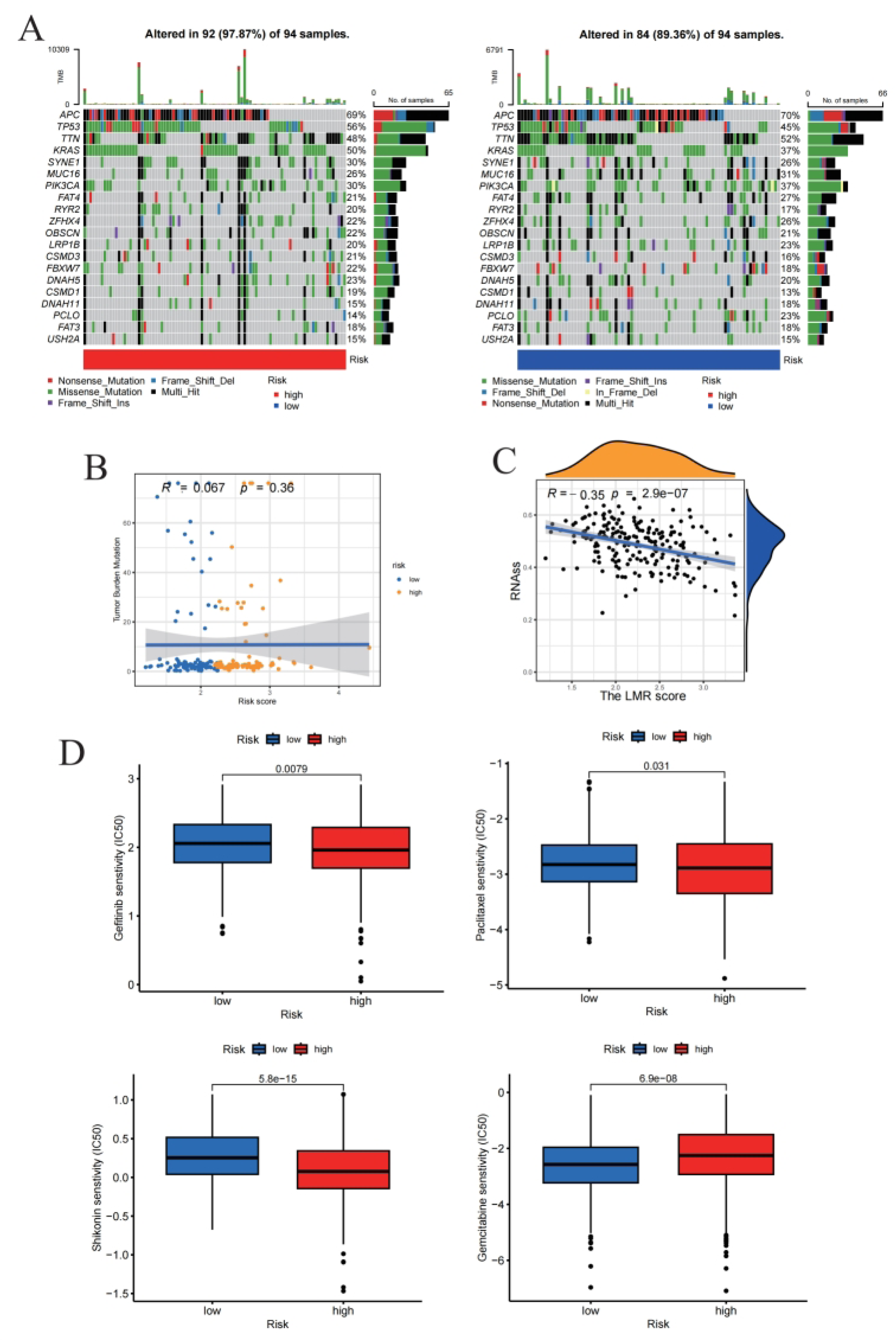
(A) The waterfall plots demonstrated gene mutation status in the low and high risk groups. The plots depicted the correlation between the LMR score and TMB (B) and CSC (D). (E) The box plots displayed the differences of chemotherapeutic sensitivity between the high and low risk groups.

## Discussion

The tumor microenvironment (TME) influences CRC development and metastasis(18). The scRNA-seq technology has been used to provide a new perspective for exploring the TME. In the present study, we characterized the primary CRC epithelium and CRLM epithelium and further identified LMR DEGs based on the basis of scRNA-seq data. The LMR DEGs were found to be enriched with pathways such as the PPAR signaling pathway. PPARs are the most well-studied fatty acid-activated transcription factors and include three subtypes: PPARα, PPARγ, and PPARδ (19). In oxidized tissues, PPARα and PPARδ are highly expressed and regulate genes involved in the oxidative phosphorylation (OXPHOS) process (20, 21). PPARγ, on the other hand, has a role in adipose tissue production and development (22). In addition, as key regulators of metabolism, PPARs can regulate the differentiation and expansion of a variety of immune cells, such as macrophages and T cells(23–25). Owing to the important functions of PPARs, an increasing number of studies have focused on the potential effects of PPARs on cancer. Aberrant activation of the PPAR signaling pathway is thought to be associated with the development of a variety of tumors, including CRC(26). A study based on single-cell sequencing and multiomics data revealed that the PPAR signaling pathway was strongly upregulated in the CRC epithelium compared with the normal epithelium; moreover, PPAR inhibitors inhibited the growth of CRC organoids (27). Our study revealed that the PPAR signaling pathway was significantly upregulated in the CRLM epithelium compared with the primary CRC epithelium. Our study complements the study of Wang et al.(27) by demonstrating at the cellular level that aberrant activation of the PPAR signaling pathway is closely associated not only with CRC occurrence but also with CRC metastasis.

The discovery of new biomarkers could improve the prognosis of CRC patients. A 15-gene risk scoring system, the LMR score, was constructed in the present study. We compared the LMR score to the AJCC stage and to the CRG score and the PRG score, which are composed of similar datasets, and found that the LMR score had satisfactory predictive power for patient DFS. In addition, the LMR score was negatively correlated with MSI status, and the high-risk score group had a higher proportion of RAS/BRAF mutations. MSI-H status is generally associated with more favorable outcomes and better responses to immune checkpoint inhibitors(28, 29), whereas RAS/BRAF mutations often indicate poorer clinical outcomes(16, 17). This molecular context aligns with the observed differential therapeutic responses: the low-risk group exhibited a significantly higher response rate to immunotherapy. Furthermore, regarding chemotherapeutic agent sensitivity, the low-risk group demonstrated greater sensitivity to gemcitabine (lower IC50), while the high-risk group showed lower IC50 values for gefitinib, paclitaxel, and shikonin. These findings indicate that the LMR score not only carries prognostic value but may also potentially inform therapeutic strategy by identifying patient subgroups more likely to benefit from immunotherapy or specific chemotherapy regimens.

We observed that the absolute value of the coefficient for *DCBLD2* was the largest for the LMR score. *DCBLD2* was first cloned from lung cancer cells with high metastatic potential and was found to be significantly overexpressed in highly metastatic lung cancer cell lines (30). *DCBLD2* was subsequently found to be significantly upregulated in a variety of tumors other than lung cancer and was associated with poor patient prognosis(31). A pan-cancer analysis based on the TCGA database confirmed this finding and revealed that *DCBLD*2 expression was associated with the activation of epithelial-mesenchymal transition (EMT) signals (32). The overregulation of EMT is considered to be one of the key factors in cancer recurrence and metastasis(33, 34). Cellular and animal experiments by Huang et al. revealed that silencing of DCBLD2 induced apoptosis in CRC cells and inhibited CRC growth (35). In the present study, we found that *DCBLD2* expression has potential for predicting the prognosis of CRC patients and that patients with high *DCBLD2* expression have a poor prognosis, which is consistent with the results of previous studies. More importantly, we found that the methylation level of the promoter region of *DCBLD2* was significantly negatively correlated with expression. The methylation level of *DCBLD2* may be a valuable biomarker since abnormalities in DNA methylation precede genetic alterations such as mutations (36, 37), but this requires more in-depth studies.

Another major focus of this study was to explore the association between the LMR score and the tumor immune microenvironment (TIME). The TIME consists mainly of various types of immune cells, and different types of immune cells play different roles in the development of tumors. For example, CD8+ T cells lyse tumor cells and release cytokines that enhance cytotoxic effects(38, 39), whereas regulatory T cells (Tregs), a subpopulation of CD4+ T cells, are essential for maintaining immune homeostasis and are generally thought to suppress T-cell responses, thereby promoting immune evasion of tumors(40, 41). However, Treg infiltration has also been found to be associated with a favorable prognosis in CRC. The reason for this discrepancy may be related to differences in the cellular phenotype, gene expression, and functional activities of Tregs (42). In addition, a variety of immune cell types such as natural killer (NK) cells(43), neutrophils(44), other subtypes of CD4+ T cells(45) and plasma cells(46), play important roles in the tumor immunity process. The complex functions of immune cells and the interactions between immune cells make the role of the TIME in tumor progression difficult to evaluate, and sometimes it even has dual effects: antitumor effects and protumor effects (47). In our study, the LMR score was associated with multiple immune cell types. To measure whether a patient’s TIME was protumor or antitumor and the association between the LMR score and immune status, we evaluated the patient’s TIME using the TIDE score. The TIME, especially the infiltration of T cells, has been proven to be closely related to the efficacy of tumor immunotherapy(48, 49). Investigators of the TIDE score constructed algorithms to characterize T-cell dysfunction in other tumor cohorts by examining the correlation between the expression of each gene in a tumor modeling cohort and the level of cytotoxic T lymphocyte (CTL) infiltration and its impact on survival(50). The TIDE score is thought to be negatively correlated with immunotherapy outcome. We found that patients with high LMR scores had higher TIDE scores and lower immunotherapy response rates, which revealed a strong association between the LMR score and TIME and provides a basis for immunotherapy for CRC patients based on the basis of the LMR score. Certainly, more clinical evidence is needed.

## Limitations

Overall, our study provides new insights into the occurrence of CRLM, as well as a reliable biomarker for the prognosis of CRC. However, there are some shortcomings in the current study. First, the interactions of various cells in the TME need to be further explored. Second, the functions of some of the genes used to constitute the LMR score have not been elucidated. Finally, the ability of the LMR score to predict DFS in patients with CRC needs to be validated in a larger cohort.

## Conclusion

In this study, we depicted the cellular landscape of CRLM and primary CRC on the basis of a scRNA-seq dataset and then identified LMR DEGs between primary CRC and CRLM at the cellular level. Based on the LMR DEGs, we constructed a risk score (the LMR score) consisting of the expression of 15 genes. The LMR score has favorable predictive power for DFS in patients with CRC. Further analysis revealed that the LMR score was significantly associated with immune infiltration, chemotherapeutic drug sensitivity, and other factors influencing the prognosis of CRC patients.

## Declaration

### Ethical approval and Accordance Statement

The datasets used in this study are publicly accessible via open repositories.

### Consent to publish

Not applicable.

### Consent to participate

Not applicable.

### Accordance Statemen

The datasets used in this study are publicly accessible via open repositories.

### Data Availability Statement

Data for this study can be found in The Cancer Genome Atlas (TCGA, https://portal.gdc.cancer.gov/) and the Gene Expression Omnibus (GEO, https://www.ncbi.nlm.nih.gov/geo/). The single-cell RNA-seq dataset analyzed in this study is available in the GEO repository under accession number GSE178318. Bulk RNA-seq data analyzed in this study were obtained from public repositories, including the GEO datasets under accession numbers GSE17536, GSE17537, GSE38832, GSE39582, and GSE161158, as well as the colorectal cancer transcriptomic, clinical, and methylation data from the TCGA projects TCGA-COAD and TCGA-READ. Mutation data were analyzed via the cBioPortal platform (https://www.cbioportal.org) using the TCGA Colorectal Adenocarcinoma (TCGA, PanCancer Atlas) dataset. Analyses of tumor immune dysfunction and exclusion were performed using the Tumor Immune Dysfunction and Exclusion (TIDE) resource (http://tide.dfci.harvard.edu).

## Acknowledgements

We are indebted to all the patients who participated in studies of TCGA and GEO databases.

## Author Contributions

Data collection and filtering: XZ, WC, RH; data analysis and figures preparation: XZ; collation of results: WC, LL, JL; article writing: XZ, HL; Review and modify article format: YX, WZ; conceived and designed this study: YX, XZ, YL. All authors read and approved the final manuscript.

